# Design and characterization of HIV-1 vaccine candidates to elicit antibodies targeting multiple epitopes

**DOI:** 10.1101/2025.01.08.632013

**Authors:** Harry B. Gristick, Harald Hartweger, Yoshiaki Nishimura, Edem Gavor, Kaito Nagashima, Nicholas S. Koranda, Priyanthi N.P. Gnanapragasam, Leesa M. Kakutani, Luisa Segovia, Olivia Donau, Jennifer R. Keeffe, Anthony P. West, Malcolm A. Martin, Michel C. Nussenzweig, Pamela J. Bjorkman

## Abstract

A primary goal in the development of an AIDS vaccine is the elicitation of broadly neutralizing antibodies (bNAbs) that protect against diverse HIV-1 strains. To this aim, germline-targeting immunogens have been developed to activate bNAb precursors and initiate the induction of bNAbs. While most pre-clinical germline-targeting HIV-1 vaccine candidates only include a single bNAb precursor epitope, an effective HIV-1 vaccine will likely require bNAbs that target multiple epitopes on Env. Here, we report a newly designed germline-targeting Env SOSIP trimer, named 3nv.2, that presents three bNAb epitopes on Env: the CD4bs, V3, and V2 epitopes. 3nv.2 forms a stable trimeric Env and binds to bNAb precursors from each of the desired epitopes. Immunization experiments in rhesus macaques and mice demonstrate 3nv.2 elicits the combined effects of its parent immunogens. Our results provide proof-of-concept for using a germline-targeting immunogen presenting three or more bNAb epitopes and a framework to develop improved next-generation HIV-1 vaccine candidates.

## Introduction

There are nearly 40 million people currently infected with HIV-1 and 1-2 million new infections each year, but only ∼77% of people living with HIV-1 have access to anti-retroviral drugs (www.who.org). An effective vaccine remains the best option to prevent new infections worldwide but has proven difficult due to factors including (*i*) the extensive genetic diversity of circulating strains, (*ii*) the fact HIV-1 is a retrovirus and once integrated, can only be cleared in rare circumstances(Allers et al., 2011), and (*iii*) unusual properties of HIV bNAbs including long CDRH3s(Walker et al., 2009, Bonsignori et al., 2011, Doria-Rose et al., 2014, Sok et al., 2014) and rare germline gene usage(McGuire et al., 2013, Jardine et al., 2013). Although most people living with HIV-1 (PLWH) generate only strain-specific or non-neutralizing antibodies (Abs), an estimated 5-20% of PLWH produce broadly neutralizing Abs (bNAbs) that neutralize a wide array of strains at low concentrations (<50 µg/mL) after being infected for several years(McCoy and Burton, 2017). These bNAbs can protect rhesus macaques (RMs) from challenge from simian HIV-1 (SHIV) infection(Gautam et al., 2016, Shingai et al., 2014), suggesting a vaccination regimen that elicits bNAbs at neutralizing concentrations would be protective(Walsh and Seaman, 2021).

The HIV-1 Envelope protein (Env), a heterotrimeric membrane glycoprotein comprising gp120 and gp41 subunits found on the surface of the virion, is responsible for viral entry into host cells and is the sole antigenic target of neutralizing Abs(Wyatt and Sodroski, 1998). Structural and biochemical studies have elucidated how bNAbs recognize Env and described correlates of neutralization breadth and potency(Ward and Wilson, 2017). A native-like soluble form of the Env trimer ectodomain (SOSIP) can be produced from most Env strains(Sanders et al., 2013, Derking and Sanders, 2021). The ectodomains of SOSIP and native virus-associated Envs have similar 3D structures(Li et al., 2020), CD4-recognition properties(Li et al., 2023), and present analogous bNAb epitopes(Derking et al., 2015), making SOSIPs ideal candidates for vaccine design to elicit bNAbs(Derking and Sanders, 2021). An impediment to generating an effective HIV-1 vaccine is that many inferred germline (iGL) precursors of characterized bNAbs do not bind with detectable affinity to native Envs on circulating HIV-1 strains or their counterpart SOSIPs(Xiao et al., 2009). Therefore, Env must be modified to bind and select for bNAb precursors *in vivo* during immunization. This approach, known as germline-targeting, assumes that a given Env must have appreciable affinity to a B cell expressing a bNAb precursor receptor in order to bind and activate that B cell lineage to initiate bNAb induction(Stamatatos et al., 2017). As an example, germline targeting has been used to select and activate bNAb precursors of the VRC01-class of bNAbs that target the CD4-binding site (CD4bs) on gp120(McGuire et al., 2016, Jardine et al., 2016, Jardine et al., 2015, Jardine et al., 2013, Medina-Ramirez et al., 2017).

The majority of current HIV-1 vaccine candidates target a single bNAb precursor lineage or epitope(Saunders et al., 2019, Willis et al., 2022, Steichen et al., 2019, Steichen et al., 2016, Gristick et al., 2023, Escolano et al., 2019, Jardine et al., 2016, Jardine et al., 2013, McGuire et al., 2013). However, recent findings suggested that administering bNAbs targeting the CD4bs, V3, and V2 epitopes on HIV-1 Env represents an optimal combination to neutralize 100% of circulating viruses in sub-Saharan Africa(Mkhize et al., 2023), the site of the majority of HIV-1 infections worldwide (www.unaids.org). Thus, induction of multiple bNAb lineages and/or bNAbs targeting multiple epitopes on HIV-1 Env is likely required to generate a protective HIV-1 vaccine that is effective worldwide. Towards this aim, we engineered immunogens based on Env SOSIP trimers that present two different epitopes designed to elicit bNAb lineages: RC1, a V3-glycan patch immunogen that elicited Abs targeting the conserved V3 epitope in animal models with a polyclonal Ab repertoire(Escolano et al., 2019), and IGT2, which targets CD4bs Abs and elicited heterologous serum neutralization in transgenic and wildtype (wt) mice(Gristick et al., 2023). A separate study modified BG505 to bind V2 bNAb precursors(Medina-Ramirez et al., 2017). Eliciting three (or more) classes of HIV-1 bNAbs could be favorable for generating an effective HIV-1 vaccine.

Although targeting of >1 bNAb epitope could be accomplished by immunizing with multiple immunogens, each including a single epitope, the simultaneous immunization of multiple designed immunogens would not prevent the immune system from making distracting Abs against the parts of each of the immunogens that were not modified for inducing Ab recognition. In contrast, a single immunogen with multiple engineered epitopes would reduce manufacturing complexity as well as display fewer off-target epitopes for distracting Abs to bind and thus provide a higher likelihood of engaging bNAb precursors (Fig. 1). Here, we describe the design and characterization of individual immunogens that elicit Abs targeting more than one epitope. The top vaccine candidate, 3nv.2, forms a stable trimeric Env that binds to three different classes of precursors of bNAbs recognizing the CD4bs, V3, and V2 epitopes (Fig. 2A). Importantly, immunization regimens using 3nv.2 elicited the combined effects of the counterpart single epitope immunogens. These experiments represent proof-of-concept results suggesting that presenting multiple bNAb epitopes on HIV-1 Env would be favorable over the standard approach of presenting a single bNAb epitope.

**Fig. 1.**
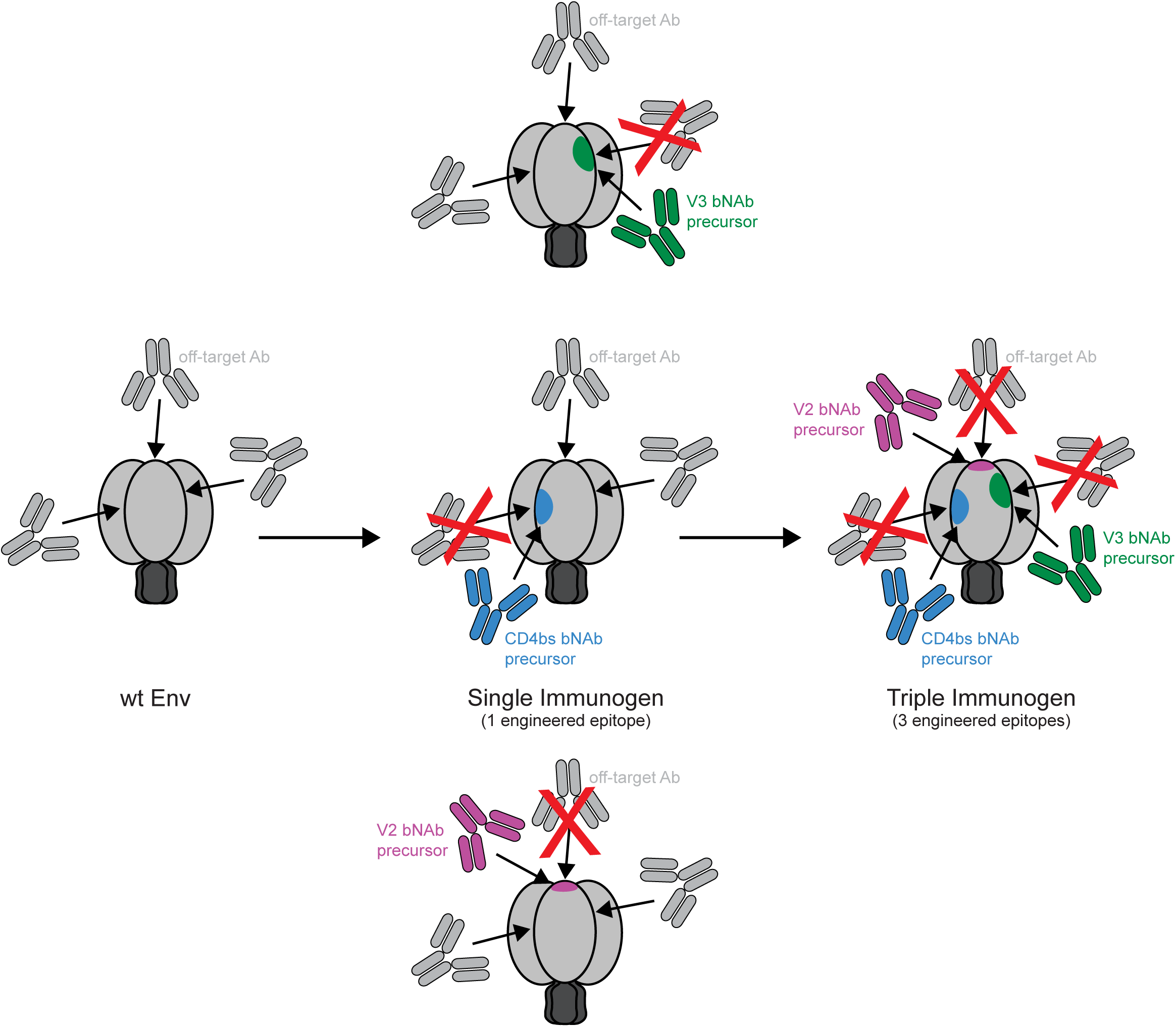
Schematic illustrating how a combined triple immunogen could elicit a lower proportion of off-target Abs than a combination of three single immunogens, each presenting only one epitope.

**Fig. 2.**
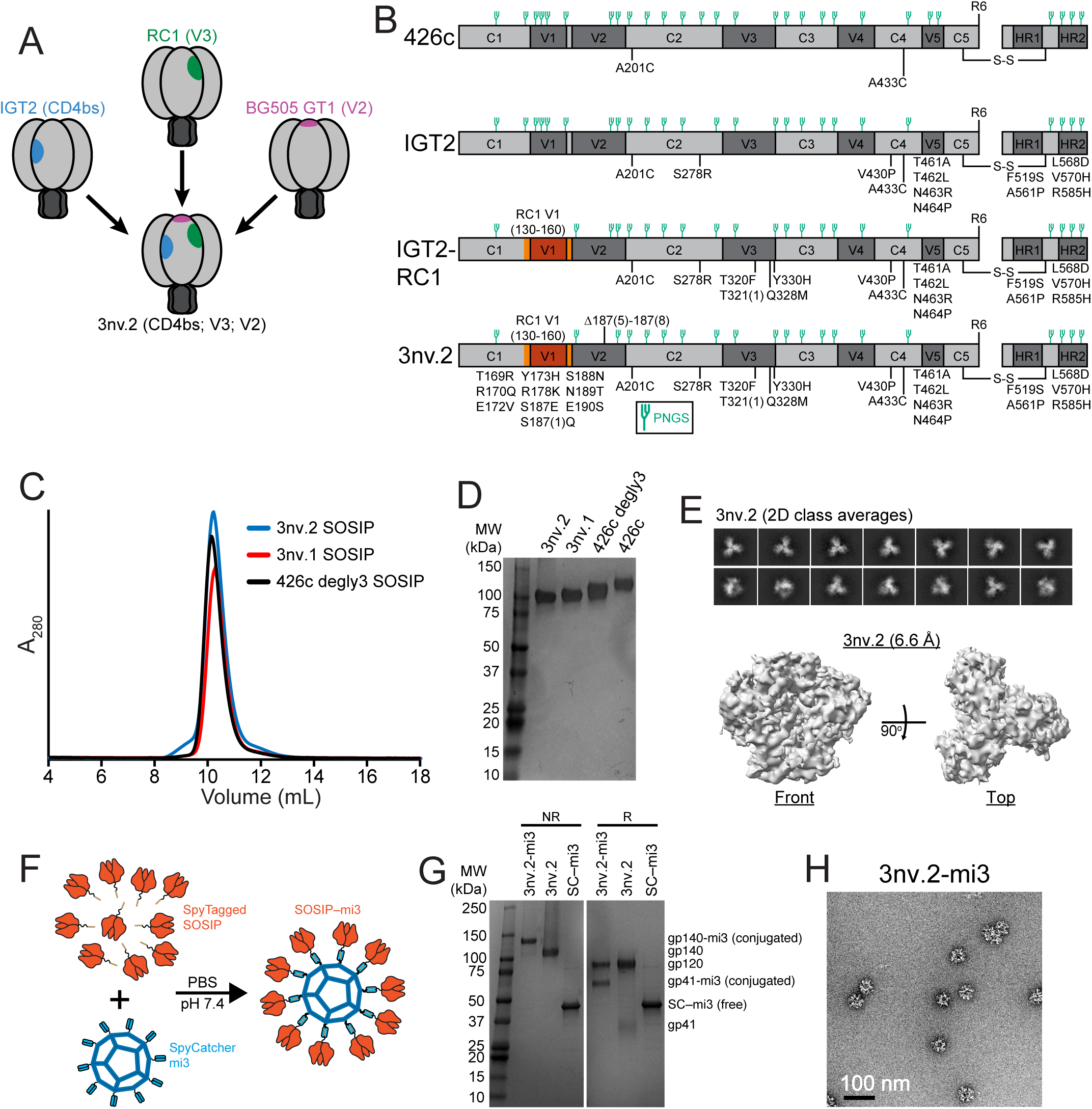
Design and biochemical characterization of 3nv.2 SOSIP. (A) Design of 426c-based triple immunogen to present CD4bs (blue), V3 (green), and V2 (purple) epitopes. (B) Schematics of single (IGT2), double (IGT2-RC1), and triple (3nv.2) immunogen constructs used in this study. (C,D) 3nv.2 and 3nv.1 triple immunogen characterization by (C) SEC and (D) SDS-PAGE. (E) Top, 2D class averages demonstrating 3nv.2 is predominantly trimeric. Bottom, 6.6 Å single-particle 3nv.2 cryo-EM density map. (F) Schematic for the generation of SOSIP-mi3 nanoparticles using the SpyCatcher-SpyTag system. (G) Characterization of purified SOSIP-mi3 nanoparticles by SDS-PAGE. R, Reduced; NR, Non-reduced; SC, SpyCatcher. (H) Negative-stain EM of SOSIP-mi3 nanoparticles. Scale bar = 100 nm.

## Results

### 3nv.2 was designed to bind to bNAb precursors targeting three Env epitopes

The design and analysis of BG505-GT1.1, an immunogen demonstrated to bind to bNAb precursors targeting the CD4bs and V2 epitopes, was previously described(Medina-Ramirez et al., 2017). Our goal was to build on these studies and engineer a priming immunogen with enhanced affinity compared to unmodified SOSIP Envs for iGL precursors of bNAbs targeting three distinct sites on Env: the CD4bs, V3, and V2 epitopes (Fig. 2A). Starting with IGT2, a clade C 426c-based SOSIP Env trimer that elicits Abs to the CD4bs(Gristick et al., 2023) (Fig. 2B), we first incorporated substitutions that enhanced targeting to the V3 epitope. This involved two distinct modifications: (1) transplanting the V2 cassette (residues 130_gp120_-160_gp120_) from RC1, a V3-targeting SOSIP immunogen(Escolano et al., 2019), into IGT2, and (2) incorporating V3-targeting residues (T320F_gp120_, T321(1)_gp120_, Q328M_gp120_, Y330H_gp120_) from RC1 into IGT2 (Fig. 2B). These combined substitutions created a CD4bs/V3-targeting double immunogen, known as IGT2–RC1 (Fig. 2B). We further modified IGT2–RC1 to include known V2 iGL-targeting residues (T169R, R170Q, E172V, Y173H, R178K, S188N, N189T, T190S), which was shown to increase binding affinity to the iGLs of V2 bNAbs including PG9 and PG16 bNAbs(Medina-Ramirez et al., 2017) (Fig. 2B). We then deleted four residues within V2 (ΔNSNK; residues 187(7)_gp120_-189_gp120_) and introduced two additional substitutions (S187E, S187(1)Q) to further enhance binding to V2 iGLs(Medina-Ramirez et al., 2017), thereby creating 3nv.2 (Fig. 2A,B). Finally, we retained five mutations (F519S, A561P, L568D, V570H, R585H) in gp41 that were demonstrated to enhance stability and increase expression levels of Env(Steichen et al., 2016) and two cysteine substitutions at residues A201_gp120_ and A433_gp120_ that form disulfide bonds to stabilize the closed conformation of Env(Joyce et al., 2017).

To create a potential boosting immunogen to be used in conjunction with a 3nv.2 prime, we started with 3nv.2 and replaced the CD4bs, V3, and V2 targeting mutations with substitutions that were more native-like and/or predicted to have lower affinity to the iGLs of interest. First, we introduced the CD4bs substitutions from IGT1, an immunogen shown to boost CD4bs responses in animal models primed with IGT2(Gristick et al., 2023). Next, we reintroduced the N156_gp120_ potential N-linked glycosylation site (PNGS) that is present in the V3-targeting immunogen 11MUTB(Steichen et al., 2016), previously shown to boost responses directed toward the V3-glycan patch when starting with the RC1 priming immunogen that lacks the N156_gp120_ PNGS(Escolano et al., 2019). Finally, we reintroduced the 4-residue deletion in the V2 loop (NSNK; residues 185e_gp120_-190_gp120_) to shepherd bNAb precursors to acquire the proper somatic hypermutations (SHMs) by introducing a more native-like environment in this region. Together, these substitutions created 3nv.1 (Fig. S1).

Both 3nv.2 and 3nv.1 triple immunogens were well-behaved in solution, monodisperse by size-exclusion chromatography (SEC), and existed as a single species in SDS-PAGE, similar to both the starting 426c SOSIP Env and the 426c degly3 variant(Borst et al., 2018) lacking PNGSs at N276_gp120_, N460_gp120_, and N463_gp120_ (Fig. 2B,C,D, Fig. S1). To assess the impact of the substitutions within 3nv.2 on the trimer structure, we solved a 6.6 Å single-particle cryo-electron microscopy (cryo-EM) structure of an untagged 3nv.2 Env expressed in Expi293 cells. Consistent with SEC and SDS-PAGE, 3nv.2 was monodisperse and predominantly trimeric as evidenced within both the 2D class averages (Fig. 2E, top) and the 3D electron density map (Fig. 2E, bottom). To enhance antigenicity and immunogenicity through avidity effects from multimerization(López-Sagaseta et al., 2016, Slifka and Amanna, 2019), we used the SpyCatcher-SpyTag system(Keeble et al., 2019, Bruun et al., 2018, Brune et al., 2016) to covalently link SpyTagged SOSIP immunogens to the 60-mer nanoparticle SpyCatcher003-mi3(Keeble et al., 2019), as we previously described for other SOSIP Env trimers(Gristick et al., 2023, Escolano et al., 2021, Escolano et al., 2019) (Fig. 2F). Efficient covalent coupling of the immunogens to SpyCatcher003-mi3 was demonstrated by SDS-PAGE (Fig. 2G), and negative-stain electron microscopy (nsEM) showed that nanoparticles were uniform in size and shape and densely conjugated with SOSIP Envs (Fig. 2H).

### 3nv.2 SOSIP trimers bind multiple bNAb precursors

To determine whether the engineered immunogens bind the desired iGL precursors, we multimerized our immunogens on mi3 nanoparticles(Keeble et al., 2019) and evaluated binding to iGL versions of IOMA(Gristick et al., 2016, Gristick et al., 2023) (CD4bs), PGT121/10-1074(Escolano et al., 2016) (V3), PG9(Medina-Ramirez et al., 2017, Sliepen et al., 2015) (V2), and PG16(Medina-Ramirez et al., 2017, Sliepen et al., 2015) (V2) using a surface plasmon resonance (SPR)-based assay. As expected, the parent immunogen, IGT2, bound to IOMA iGL (CD4bs) but not to PGT121/10-1074 iGL (V3), PG9 iGL (V2), or PG16 iGL (V2) (Fig. 3A). The dual immunogen IGT2-RC1 bound to both IOMA iGL (CD4bs) and PGT121/10-1074 iGL (V3), but not PG9 or PG16 iGLs (V2) (Fig. 3A).

**Fig. 3.**
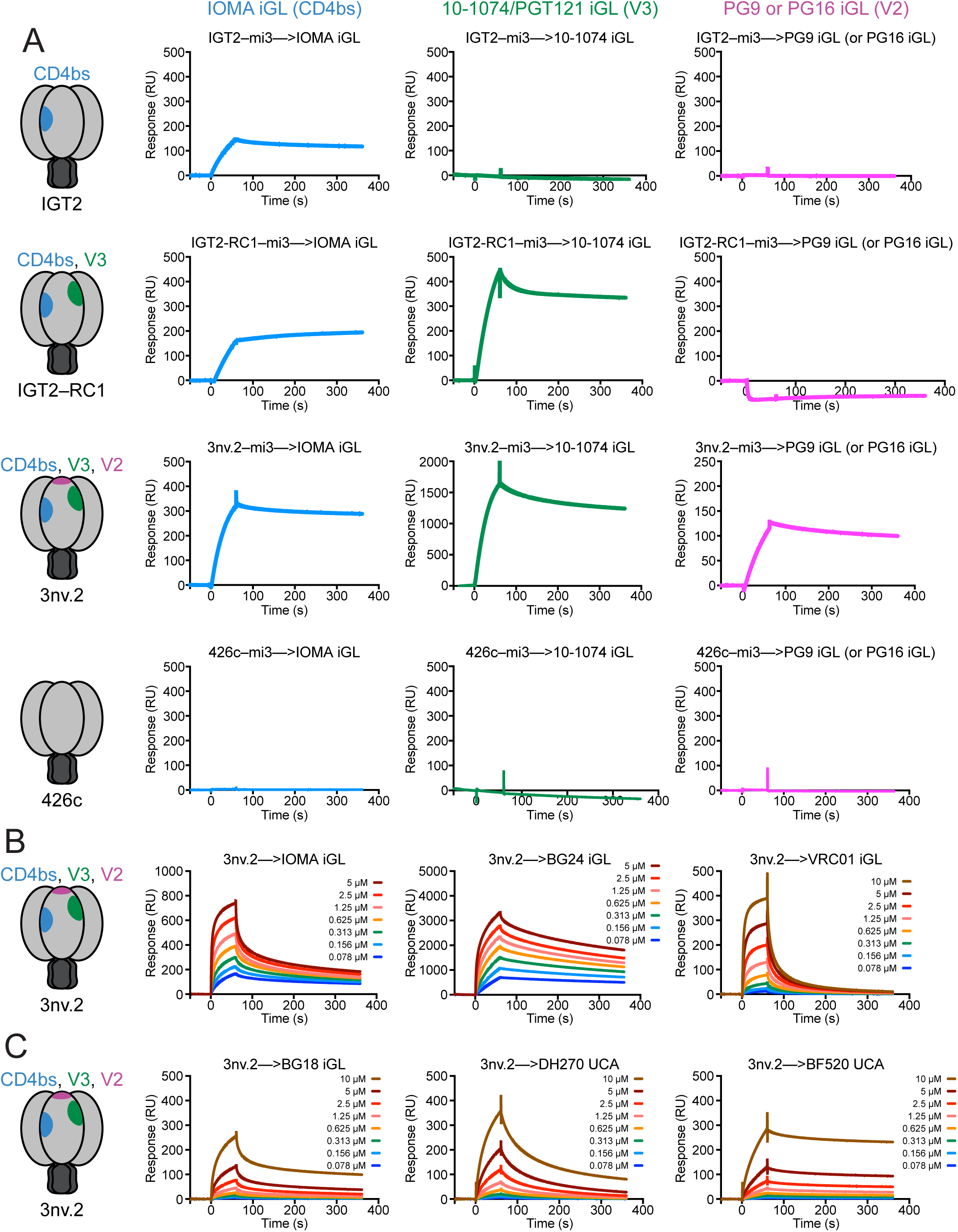
3nv.2 binds to iGLs targeting three bNAb epitopes. (A) SPR sensorgrams of SOSIP-mi3 nanoparticles injected over bNAb iGLs at a concentration of ∼0.5mg/mL. First row: the CD4bs-specific single immunogen IGT2 only binds to CD4bs bNAb precursors (IOMA iGL, left), and not V3 (10-1074/PGT121 iGL, middle) or V2 (PG9 or PG16 iGL, right) bNAb precursors. Second row: incorporating V3-targeting mutations into IGT2 creates an immunogen that binds CD4bs and V3 bNAb precursors, but not V2 bNAb precursors. Third row: incorporating V3- and V2-targeting residues into the CD4bs-targeting IGT2 SOSIP creates an immunogen (3nv.2) that binds to the CD4bs (IOMA iGL, left), V3 (10-1074/PGT121 iGL, middle), and V2 (PG9 or PG16 iGL, right) bNAb precursors. Fourth row: the parental 426c Env does not bind to any of the bNAb precursors. (B) 3nv.2 SOSIP injected in dilution series over bNAb iGLs starting at top concentrations of 10 µM or 5 µM as indicated. 3nv.2 SOSIP binds to multiple CD4bs precursors including IOMA iGL (left), BG24 iGL (middle), and VRC01 iGL (right). (C) 3nv.2 SOSIP injected in dilution series over bNAb UCAs and an iGL starting at a top concentration of 10 µM. 3nv.2 SOSIP binds to multiple V3 precursors including BG18 iGL (left), DH270 UCA (middle), and BF520 UCA (right). Representative sensorgrams are from at least two independent experiments.

However, the triple immunogen 3nv.2 bound to iGLs from all three classes - IOMA iGL (CD4bs), PGT121/10-1074 iGL (V3), and PG9 or PG16 iGLs (V2) (Fig. 3A). Importantly, 3nv.2 did not exhibit reduced binding for IOMA iGL compared to IGT2, demonstrating that modifying the V3 and V2 epitopes had no effect on the antigenicity of the CD4bs. As expected due to avidity effects, the binding interaction of the multimerized immunogens exhibited a slow off-rate that produced a strong association between 3nv.2 and the iGL IgGs (Fig. 3A). Importantly, the observed binding was due to the germline-targeting mutations introduced into our immunogens and not only due to increased avidity effects, as we observed no binding of the wt 426c-mi3 nanoparticles to any of the iGL IgGs (Fig. 3A). While 3nv.2 was designed to bind to IOMA iGL, 3nv.2 also bound to additional CD4bs precursors of different classes including BG24iGL and VRC01 iGL (Fig. 3B). Similarly, 3nv.2 not only bound to PGT121/10-1074 iGL, but also to additional V3 bNAb precursors within the V3 epitope supersite such as BG18 iGL(Freund et al., 2017, Barnes et al., 2018, Steichen et al., 2019), and the unmutated common ancestors (UCAs) of DH270(Bonsignori et al., 2017a) and BF520(Simonich et al., 2016) (Fig. 3C). In summary, 3nv.2 targets a diverse set of bNAb precursors in individual epitopes and is the first reported HIV-1 immunogen successfully engineered to bind bNAb precursors presenting three different Env epitopes.

### Animal immunizations with 3nv.2

To evaluate a 3nv.2-based immunization regimen in animals with a polyclonal Ab repertoire, we primed 15 rhesus macaques (RMs) with either 3nv.2-mi3 (*n* = 5), IGT2-mi3(Gristick et al., 2023) (*n* = 5), or RC1-mi3(Escolano et al., 2021, Escolano et al., 2019) (*n* = 5), followed in each case by sequential immunization with a related boosting antigen (3nv.1-mi3, IGT1-mi3, or 11MUTB-mi3, respectively) (Fig. 4A). Serum from animals primed with IGT2-mi3 and boosted with IGT1-mi3 only neutralized IGT2- and IGT1-based pseudoviruses (Fig. 4B,C). However, animals primed with 3nv.2-mi3 and boosted with 3nv.1-mi3 exhibited the combined effects of both the IGT2/IGT1 and RC1/11MUTB immunization regimens and displayed potent neutralization of pseudoviruses generated from the IGT2, IGT1, RC1, and 11MUTB immunogens (Fig. 4B-E). Importantly, the 426c-based (clade C) 3nv.2/3nv.1–mi3 immunization regimen elicited heterologous neutralization against RC1 and 11MUTB pseudoviruses, which were derived from BG505-based (clade A) Envs(Escolano et al., 2021). Consistent with our previous results for an IGT2-based regimen(Gristick et al., 2023), priming with IGT2-mi3 followed by boosting with IGT1-mi3 elicited strong serum binding responses that were CD4bs-specific, as demonstrated by ELISA with IGT1 and IGT1 CD4bs knockout (KO) proteins(Gristick et al., 2023) (Fig. 4F). Notably, however, serum binding responses were significantly more CD4bs-specific in animals that received the 3nv triple immunogens (0.001 < p ≤ 0.01) (Fig. 4F).

**Fig. 4.**
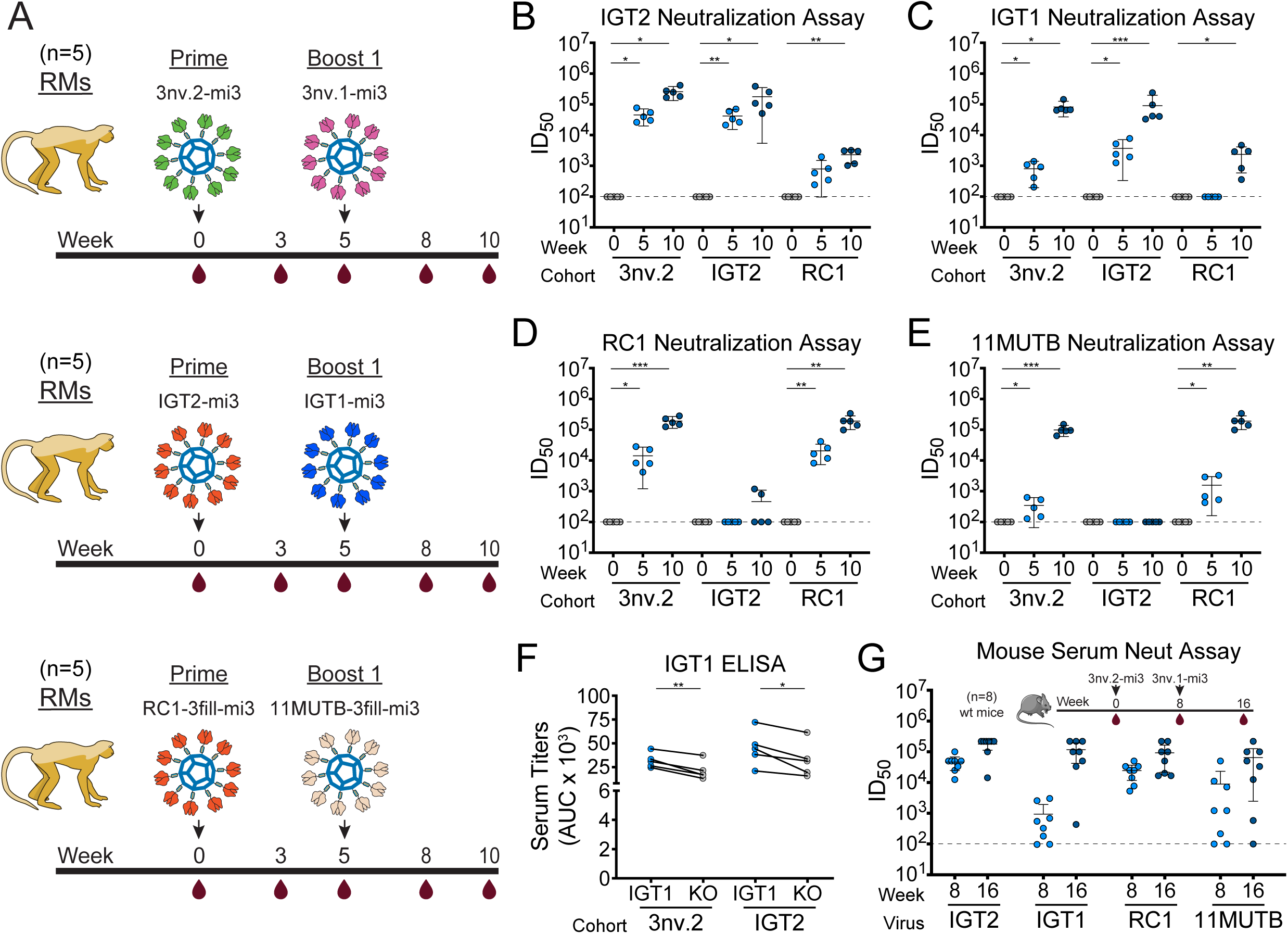
3nv.2 triple immunogen elicits combined responses of IGT2 and RC1 immunogens in rhesus macaques. (A) Schematic and timeline of immunization regimens for rhesus macaques (*n* = 5 per group). (B-E) Serum neutralization ID50s against (B) IGT2, (C) IGT1, (D) RC1, and (E) 11MUTB pseudoviruses. The dotted line at y = 10^2^ indicates the lowest dilution evaluated. Significance was demonstrated using an unpaired t test (p ≤ 0.05). (F) Serum ELISA binding to IGT1 and IGT1 CD4bs KO SOSIPs for RMs immunized with IGT2-mi3 or 3nV.2-mi3. All samples are from week 8 post prime immunizations. Significance was demonstrated using a paired t test (p ≤ 0.05). (G) Mouse serum neutralization ID50s against IGT2, IGT1, RC1, and 11MUTB pseudoviruses (*n* = 8). The dotted line at y = 10^2^ indicates the lowest dilution evaluated.

We also immunized wt C57BL/6J mice with 3nv.2-mi3 and 3nv.1-mi3 (Fig. 4G). As in RMs, serum from immunized mice also neutralized the IGT2, IGT1, RC1, and 11MUTB pseudoviruses, demonstrating the 3nv.2-mi3/3nv.1-mi3 regimen elicits cross-clade neutralizing activity in two animal models (Fig. 4G).

## Discussion

Current germline-targeting HIV-1 vaccine candidates generally target a single bNAb precursor lineage or epitope. However, the induction of multiple bNAb lineages targeting multiple epitope supersites on HIV-1 Env is likely required to generate a protective HIV-1 vaccine. Although this could be accomplished by immunizing with multiple immunogens, each presenting a different epitope, we hypothesize that simultaneous immunization with designed immunogens would not prevent the immune system from making distracting Abs against the parts of each of the immunogens that are not targeted for recognition. In other words, in a single immunogen with germline-targeting mutations in 3 individual epitopes, each Env presents a higher proportion of desired Ab epitopes than are presented in three individual one-target immunogens (Fig. 1). Here, we describe results that provide a framework to develop improved next-generation HIV-1 immunogens that present three or more epitopes.

Our top candidate triple immunogen, 3nv.2, forms a stable trimeric Env and elicits the combined effects of its parent immunogens, IGT2(Gristick et al., 2023) and RC1(Escolano et al., 2021, Escolano et al., 2019), thereby providing a platform that can be modified to present additional epitopes outside of the CD4bs, V3, and V2 regions in order to improve the priming immunization to select a large and diverse set of bNAb precursors. One such additional target is the fusion peptide (FP) epitope, found near the base of the trimer(Lee et al., 2016, Kong et al., 2016). Enhancing FP bNAb-directed responses is possible with limited Env modifications, which can be accomplished by deleting glycans surrounding the FP (e.g., N611_gp41_ PNGS)(Wang et al., 2024, Kong et al., 2019).

In addition to modifying epitopes outside of the CD4bs/V3/V2 regions, 3nv.2 could also be modified to target an even more diverse panel of bNAb precursors for each of the three epitopes it currently presents. For example, the CD4bs could be modified to also bind 8ANC131-class (VH1-46 derived)(Zhou et al., 2015) and CDRH3-dominated classes of CD4bs Abs in addition to the VRC01-class (VH1-2 derived) and IOMA-class bNAbs that 3nv.2 already binds(Gristick et al., 2023). Similarly, the V3 loop could be modified to bind additional V3-glycan patch precursors with different binding modes compared to its RC1 precursor, which targets PGT121/10-1074 precursors(Escolano et al., 2019), such as the iGL of EPCT112(Molinos-Albert et al., 2023), and the V2 loop could be designed to additionally select for targets such as PGDM1400 iGL(Willis et al., 2022). As with 3nv.2, this could be done using rational design, or in the type of high-throughput display library screen that was used to select IGT1 and IGT2(Gristick et al., 2023).

Another method to improve upon our results and elicit a more diverse bNAb response is to generate a panel of 3nv Envs from diverse HIV-1 strains, each containing three or more engineered epitopes per trimer. Analogous to experiments using mosaic nanoparticles decorated with receptor-binding domains from different sarbecoviruses(Cohen et al., 2024, Cohen et al., 2022, Cohen et al., 2021), germline-targeting mutations from 3nv.2 could be introduced into Envs from different HIV-1 clades to attempt to immunofocus responses to the desired engineered bNAb epitopes.

Although it was previously shown that engineered priming immunogens can successfully select and expand the desired bNAb precursors(Gristick et al., 2023, Saunders et al., 2019, Jardine et al., 2015, Escolano et al., 2019, Caniels et al., 2023), an effective boosting regimen has not yet been developed to efficiently shepherd those initial responses into bNAbs. In the immunization studies reported here, 3nv.2 and 3nv.1 appear to be priming epitope-specific responses; however, more experiments will be required to identify an appropriate boosting regimen to elicit NAbs with the breadth and potency required for an effective HIV-1 vaccine. Although high-throughput display methods have been applied to select priming immunogens with high affinity to the germline forms of bNAbs(Gristick et al., 2023, Steichen et al., 2019, Jardine et al., 2013), boosting immunogens have been selected in a low throughput and mostly empirical manner(Escolano et al., 2021, Chen et al., 2021, Gristick et al., 2023, Caniels et al., 2023, Zhang et al., 2021). A potential way to identify appropriate boosting immunogens is to examine Ab-virus co-evolution(Bonsignori et al., 2017b) using a SHIV infection model(Li et al., 2016). Recent studies demonstrated that SHIV-infected rhesus macaques develop HIV-1 bNAbs by means of Env-Ab coevolutionary pathways that recapitulate those that occur in HIV-1 infected humans(Roark et al., 2021). Importantly, Abs targeting the CD4bs, V3, V2, and FP epitopes similar to those isolated from individuals living with HIV-1 have been elicited in SHIV-infected RMs(Roark et al., 2021, Wang et al., 2024). Thus, Env sequences selected during bNAb development in SHIV-infected RMs could be exploited and used as boosting immunogens to elicit bNAbs in an immunization regimen. For example, top candidate combined immunogens such as 3nv.2 could be used in a SHIV infection model to identify better boosting immunogens specific for multiple bNAb epitopes that have a higher likelihood of shepherding initial primed responses to develop into bNAbs.

## Methods

### Ab, gp120, and Env trimer expression and purification

Env immunogens were expressed as soluble SOSIP.664 native-like Env trimers(Sanders et al., 2013) by transient transfection in Human Embryonic Kidney (HEK) 293-6E cells (National Research Council of Canada) or Expi293 cells (Thermo Fisher Scientific) as described(Gristick et al., 2023). For SpyTagged SOSIPs, a 16-residue SpyTag003 sequence(Keeble et al., 2019) was added to the C-terminus. Proteins were expressed. Proteins were purified from transfected cell supernatants by 2G12 affinity chromatography followed by SEC purification using a 10/300 or 16/600 Superdex 200 (GE Healthcare) column equilibrated in 20 mM Tris (pH 8.0), 150 mM NaCl (TBS) for untagged versions or 20 mM sodium phosphate (pH 7.5) and 150 mM NaCl (PBS) for SpyTagged versions as described(Gristick et al., 2023). Soluble Envs were stored at 4°C in TBS for untagged versions or PBS for SpyTagged versions.

The iGL sequences of IOMA, BG24, VRC01, 10-1074/PGT121, BG18, PG9 and PG16 IgGs were derived as previously described(Jardine et al., 2013, Steichen et al., 2019, Gristick et al., 2023, Escolano et al., 2016, Sliepen et al., 2015, Freund et al., 2017, Barnes et al., 2018, Dam et al., 2022a). The UCA IgGs sequences of DH270 and BF520 were derived as previously described(Bonsignori et al., 2017a, Simonich et al., 2016). All IgGs were expressed by transient transfection in Expi293 cells and purified from cell supernatants using MabSelect SURE (Cytiva) columns followed by SEC purification using a 10/300 or 16/600 Superdex 200 (GE Healthcare) column equilibrated in PBS(Gristick et al., 2023).

### Preparation of SOSIP-mi3 nanoparticles

SpyCatcher003-mi3 nanoparticles were prepared from BL21 (DE3)-RIPL Escherichia coli (Agilent) transformed with a pET28a SpyCatcher003-mi3 gene(Rahikainen et al., 2021) including an N-terminal 6x-His tag as described(Cohen et al., 2022, Cohen et al., 2021). Briefly, cell pellets from transformed bacteria were lysed with a cell disruptor in the presence of 2.0 mM phenylmethylsulfonyl fluoride (Sigma-Aldrich). Lysates were spun at 21,000g for 30 min and filtered with a 0.2 μm filter, and mi3 nanoparticles were isolated by ammonium sulfate precipitation followed by SEC using a HiLoad 16/600 Superdex 200 (GE Healthcare) column equilibrated with TBS. SpyCatcher003-mi3 nanoparticles were stored at 4°C and used for conjugations for up to two weeks after filtering with a 0.2 μm filter and spinning for at 14,000xg for 30 min at 4°C.

SOSIP-mi3 nanoparticles were made by incubating purified SpyCatcher003-mi3 with a 2-fold molar excess (SOSIP protomer to mi3 subunit) of purified SpyTagged SOSIP overnight at room temperature in PBS as described(Gristick et al., 2023). Conjugated SOSIP-mi3 nanoparticles were separated from free SOSIPs by SEC on a Superose 6 10/300 column (GE Healthcare) equilibrated with PBS. Fractions corresponding to conjugated mi3 nanoparticles were collected and analyzed by SDS-PAGE. Concentrations of conjugated mi3 nanoparticles were determined using the absorbance at 280 nm as measured on a Nanodrop spectrophotometer (Thermo Fisher Scientific).

### Characterization of SOSIP-mi3 nanoparticles

SOSIP-mi3 nanoparticles were characterized using ns-EM to confirm stability and SOSIP conjugations to SpyCatcher-mi3. Briefly, SOSIP-mi3 nanoparticles were diluted to 20 μg/ml in 20 mM Tris pH 8.0, 150 mM NaCl, and 4 μL of sample were applied onto freshly glow-discharged 300-mesh copper grids. Samples were incubated on a grid for 60 s, and excess sample was blotted with filter paper (Whatman). Uranyl formate (4 μL) was added for 60 s, and excess stain was then removed with filter paper. Staining with uranyl formate was repeated one more time and grids were left to air dry. Prepared grids were imaged on a 120 keV Tecnai T12 (FEI) transmission electron microscope using an Ultrascan 2k x 2k CCD (Gatan) camera at 21,000x magnification.

### SPR binding studies

SPR measurements were performed on a Biacore T200 (GE Healthcare) at 25 °C in HBS-EP+ (10mM Hepes, 150mM NaCl, 3mM EDTA, 0.005% Tween-20) (GE Healthcare) running buffer. IgGs were directly immobilized onto a CM5 chip (GE Healthcare) to ∼10,000-20,000 resonance units (RUs) using primary amine chemistry (Biacore Manual). SOSIP–mi3 samples were injected at a concentration of ∼0.5 mg/mL to verify binding to IOMA iGL IgG, 10-1074 iGL IgG, and PG9 or PG16 iGL IgG, as demonstrated in Figure 3A. Experiments were performed at least twice and representative sensorgrams are shown in Figure 3A. To assess binding of 3nv.2 to IgG forms of bNAb precursors to the CD4bs (Fig. 3B) and V3 epitopes (Fig. 3C), a concentration series of unconjugated 3nv.2 SOSIP was injected over the flow cells at increasing concentrations (top concentration 10 μM or 5 µM) at a flow rate of 60 μL/min for 60 s and allowed to dissociate for 300 s. Regeneration of flow cells was achieved by injecting one pulse of 10 mM glycine pH 3.0 at a flow rate of 90 μL/min. SPR experiments were used to qualitatively monitor binding rather than to derive binding affinities or kinetic constants, which can not be accurately determined due to avidity effects in this experimental setup(Rich and Myszka, 2010, Rich and Myszka, 2011).

### Cryo-EM sample preparation

An unliganded 3nv.2 SOSIP Env trimer structure was obtained from an epitope mapping(Turner et al., 2023) experiment in which purified 3nv.2 SOSIP was incubated overnight with polyclonal Fabs. Fab-3nv.2 SOSIP Env complexes were purified by SEC on a Superose 6 Increase 10/300 GL analytical column (GE Healthcare Life Sciences) in TBS buffer. Fractions corresponding to Fab-SOSIP complexes were concentrated to a final concentration of 2 mg/mL using a 50 kDa spin concentrator (Millipore). Immediately before deposition on grids, a 0.5% (w/v) octyl-maltoside fluorinated solution (Anatrace) was added to the protein sample to achieve a final concentration of 0.02% (w/v) as described(DeLaitsch et al., 2024). 3 μL of a protein sample was applied to freshly glow-discharged Quantifoil R1.2/1.3 grids (300 Cu mesh; Electron Microscopy Sciences), which had been treated for 1 minute at 20 mA using a PELCO easiGlow device (Ted Pella). The grid was plunge frozen using a Mark IV Vitrobot (Thermo Fisher) at 22°C and 100% humidity. Blotting was performed with Whatman No. 1 filter paper for 3 s with a blot force of 0. Finally, the sample was vitrified by rapid plunging into liquid ethane cooled by liquid nitrogen.

### Data collection and processing

Single-particle cryo-EM data acquisition was carried out on a 200 kV Talos Arctica (Thermo Fisher) microscope. Automated data collection was done using SerialEM software(Mastronarde, 2005), employing beam-image shift across a 3×3 grid of 1.2 µm holes, with one exposure per hole. 40-frame movies were captured in a super-resolution mode using a K3 camera (Gatan) with a pixel size of 0.435 Å (45,000x magnification) (Table S1). A summary of the data collection parameters is in Table S1, and the data processing workflow is in Figure S2. Data processing was carried out using cryoSPARC v.2.15(Punjani et al., 2017) (Fig. S2).

Briefly, 1200 cryo-EM movies were patch motion-corrected within cryoSPARC(Punjani et al., 2017) to account for beam-induced motion, including dose weighting, following binning of super-resolution frames. For CTF parameter estimation, non-dose-weighted micrographs were processed using the Patch CTF job in cryoSPARC(Punjani et al., 2017). Micrographs displaying poor CTF fits or evidence of crystalline ice in their power spectra were excluded from further analysis. Particle picking was performed in cryoSPARC(Punjani et al., 2017) using Blob picker for reference-free selection, and particles were extracted using the Particle Extraction Job with a box size of 360 Å. Extracted particles were subjected to several rounds of 2D classification, and the best class averages representing different views of unliganded 3nv.2 Env were used to generate two *ab initio* models. The best ab initio model was further refined to generate a final 3D volume using homogenous refinement by applying either C1 or C3 symmetry. ChimeraX (v1.8)(Pettersen et al., 2021) was used to visualize cryo-EM density maps and prepare structure figures.

### Animal immunizations and sampling

Fifteen RMs (Macaca mulatta) of Indian genetic origin were housed in a Biosafety Level 2 National Institute of Allergy and Infectious Diseases (NIAID) facility and cared for in accordance with the Guide for Care and Use of Laboratory Animals report number National Institutes of Health (NIH) 82-53 (Department of Health and Human Services, Bethesda, 1985). All RM procedures and experiments were performed according to protocols approved by the IACUC of NIAID, NIH. The RMs used in this study did not express the major histocompatibility complex class I Mamu-A*01, Mamu-B*08, and Mamu-B*17 alleles. RMs were immunized subcutaneously in the medial inner forelegs and hind legs (a total of four sites per animal) with 200 μg of the indicated SOSIP-mi3 adjuvanted in SMNP, a particulate saponin/TLR agonist vaccine adjuvant(Silva et al., 2021) (375 U per animal) as described(Escolano et al., 2021). Immunizations and blood samples were obtained from naïve and immunized macaques at time points indicated in Fig. 4A.

All mouse experiments were conducted with approval from the Institutional Review Board and the Institutional Animal Care and Use Committee (IACUC) at the Rockefeller University. C57BL/6J mice (Jackson Laboratories) were housed at a temperature of 22°C and humidity of 30 to 70% in a 12-hour light/dark cycle with ad libitum access to food and water. Male and female mice aged 8 to 13 weeks at the start of the experiment were used throughout. Sample sizes were not calculated a priori. Given the nature of the comparisons, mice were not randomized into each experimental group, and investigators were not blinded to the group allocation. Instead, experimental groups were age- and sex-matched. Mice were immunized intraperitoneally with 10 µg SOSIP-mi3 nanoparticles in 100 µL PBS with 1 U SMNP adjuvant. Serum samples were collected throughout the experiment by submandibular bleeding.

### Enzyme-linked immunosorbent assays

Serum ELISAs were performed using randomly biotinylated SOSIP trimers(Dam et al., 2022b) using the EZ-Link NHS-PEG4-Biotin Kit (Thermo Fisher Scientific) according to the manufacturer’s guidelines. Biotinylated SOSIP timers were immobilized on streptavidin-coated 96-well plates (Thermo Fisher Scientific) at a concentration of 2-5 μg/ml in TBS-T (20 mM Tris (pH 8.0), 150 mM NaCl, and 0.1% Tween 20) supplemented with 1% BSA for 1 hour at room temperature. After washing the plates in TBS-T, the plates were incubated with a threefold concentration series of rhesus macaque serum at a top dilution of 1:100 in blocking buffer for 2-3 hours at room temperature. After washing plates with TBS-T, horseradish peroxidase (HRP)-conjugated goat anti-human multispecies IgG Ab (Southern Biotech, #2014-05) was added at a dilution of 1:8000 in blocking buffer and incubated for 1 hour at room temperature. After washing the plates with TBS-T, 1-Step Ultra TMB substrate (Thermo Fisher Scientific) was added for ∼3 min. Reactions were quenched by the addition of 1 N HCl, and absorbance at 450 nm was measured using a plate reader (BioTek).

### In vitro neutralization assays

Pseudovirus neutralization assays(Montefiori, 2005) were conducted in-house as described(Gristick et al., 2023), and repeated at least twice for each reported value. Pseudoviruses made using Envs from immunogens were prepared as described(Gristick et al., 2023, Escolano et al., 2021). Briefly, RC1 and 11MUTB pseudoviruses were generated from the clade A Env BG505 and included substitutions from the RC1 and 11MUTB immunogens^39^, and IGT2 and IGT1 pseudoviruses were generated from the clade C Env 426c and included substitutions from the IGT2 and IGT1 immunogens^29^. For assessing neutralization by polyclonal Abs, serum samples were heat-inactivated at 56°C for 30 min before being added to the neutralization assays, and then neutralization was evaluated in duplicate with an eight-point, four-fold dilution series starting at a dilution of 1:20. The serum dilution responsible for 50% neutralization (ID_50_) is reported for all serum samples.

### Statistical analysis

Comparisons between groups for ELISAs and neutralization assays were calculated using an unpaired or paired t test in Prism 10.4.1 (Graphpad), as indicated. Differences were considered significant when p values were less than 0.05. p values are in relevant figures at the top of the plot, with asterisks denoting level of significance (* denotes 0.01 < p ≤ 0.05, ** denotes 0.001 < p ≤ 0.01, *** denotes 0.0001 < p ≤ 0.001, and **** denotes p ≤ 0.0001).

## Abbreviations

Ab: antibody
bNAb: broadly neutralizing Ab
CD4bs: CD4 binding site
ELISA: enzyme-linked immunosorbent assay
EM: electron microscopy
Env: envelope of HIV-1 with gp120 and gp41 subunits
FP: fusion peptide
HEK: human embryonic kidney
iGL: inferred germline
PNGS: potential N-linked glycosylation site
RM: rhesus macaque
SEC: size-exclusion chromatography
SHIV: simian HIV-1
SHM: somatic hypermutation
SOSIP: native-like soluble gp140 Env trimer
SPR: surface plasmon resonance
UCA: unmutated common ancestor
wt: wildtype

## Online supplemental material

**Fig. S1** shows the design schematics of constructs used in this study. **Fig. S2** shows the data processing of the 3nv.2 SOSIP cryo-EM dataset.

## Data and code availability

The cryo-EM map of 3nv.2 SOSIP was deposited to the Electron Microscopy Data Bank (EMDB) and has the accession code: EMD-48440. This paper does not report atomic models or original code. Additional information can be made available upon request.

## Acknowledgements

We thank D. J. Irvine and his laboratory members (MIT and Scripps Research) for providing SMNP adjuvant and Jost Vielmetter, Kaya Storm, Annie Rorick, and the Caltech Protein Expression Center in the Beckman Institute for expression assistance. Electron microscopy was performed in the Caltech Cryo-EM Center with assistance from Songye Chen. This work was supported by the National Institute of Allergy and Infectious Diseases (NIAID) Grants HIVRAD P01AI100148 (P.J.B., M.C.N.), the Bill and Melinda Gates Foundation Collaboration for AIDS Vaccine Discovery (CAVD) grant INV-002143 (P.J.B., M.A.M., and M.C.N.), and the Intramural Research Program of the NIAID (M.A.M. and Y.N.). This manuscript is the result of funding in whole or in part by the National Institutes of Health (NIH). It is subject to the NIH Public Access Policy. Through acceptance of this federal funding, NIH has been given a right to make this manuscript publicly available in PubMed Central upon the Official Date of Publication, as defined by NIH.

## Author contributions

**Harry B. Gristick:** Conceptualization, Formal Analysis, Funding Acquisition, Investigation, Methodology, Project Administration, Resources, Supervision, Validation, Visualization, Writing – Original Draft, and Writing – Review & Editing. **Harald Hartweger:** Funding Acquisition, Investigation, Methodology, Resources, Writing – Review & Editing. **Yoshiaki Nishimura:** Funding Acquisition, Investigation, Methodology, Writing – Review & Editing. **Edem Gavor:** Formal Analysis, Investigation, Methodology, Resources, Validation, Visualization, and Writing – Review & Editing. **Kaito Nagashima:** Investigation, Resources, Validation, Writing – Review & Editing. **Nicholas S. Koranda:** Formal Analysis, Investigation, Methodology, Resources, Writing – Review & Editing. **Priyanthi N.P. Gnanapragasam:** Investigation and Resources. **Leesa M. Kakutani:** Investigation and Resources. **Luisa Segovia:** Investigation and Resources. **Olivia Donau:** Investigation and Resources. **Jennifer R. Keeffe:** Formal Analysis, Funding Acquisition, Project Administration, Resources, Supervision, Validation, and Writing – Review & Editing. **Anthony P. West Jr.:** Formal Analysis, Funding Acquisition, Project Administration, Resources, Software, Supervision, Validation, and Writing – Review & Editing. **Malcolm A. Martin:** Funding Acquisition, Project Administration, Supervision, and Writing – Review & Editing. **Michel C. Nussenzweig:** Funding Acquisition, Project Administration, Supervision, and Writing – Review & Editing. **Pamela J. Bjorkman:** Conceptualization, Formal Analysis, Funding Acquisition, Methodology, Project Administration, Supervision, Validation, Visualization, Writing – Original Draft, and Writing – Review & Editing.

---

**Fig. S1.**
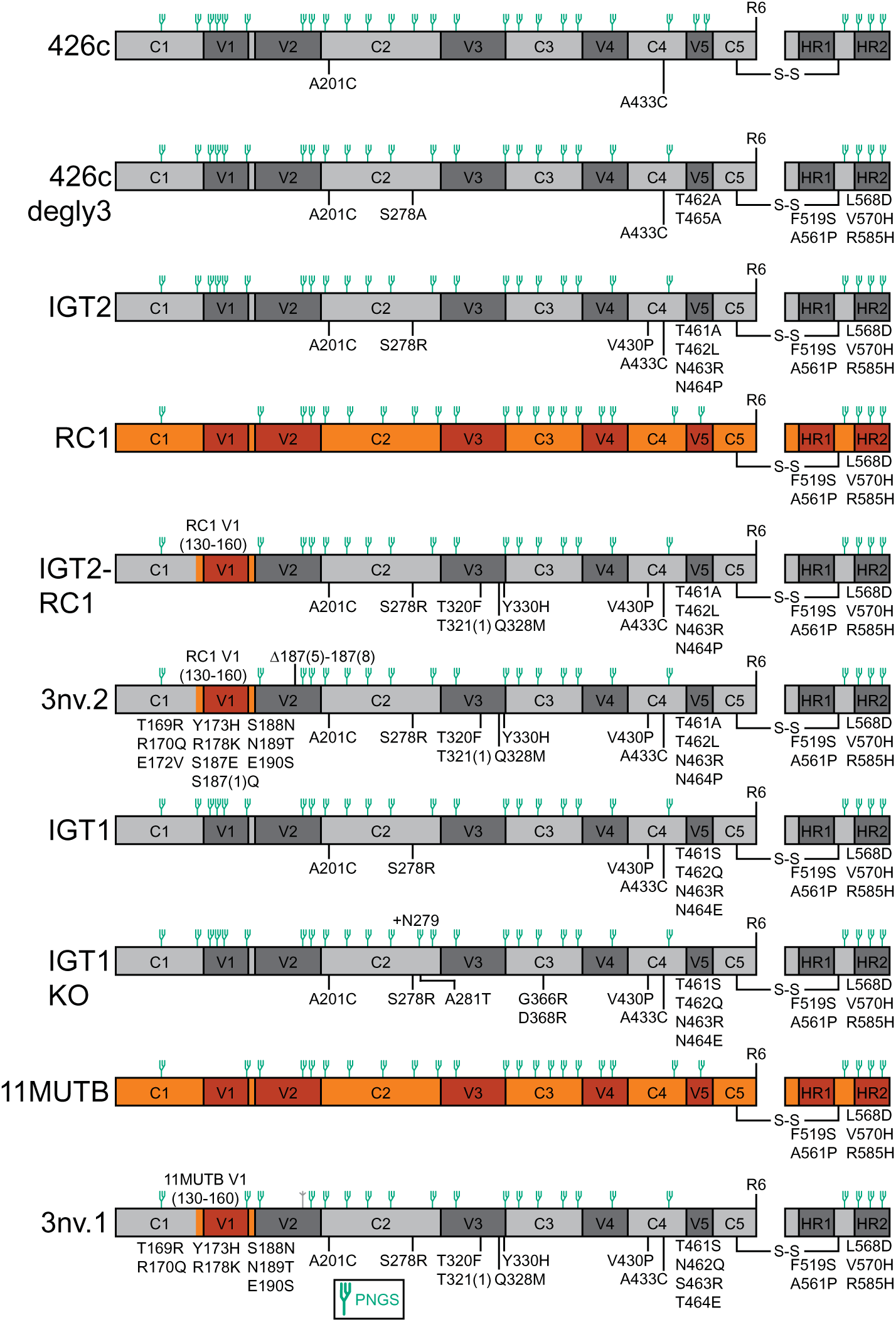
Schematics of constructs used in this study.

**Fig. S2.**
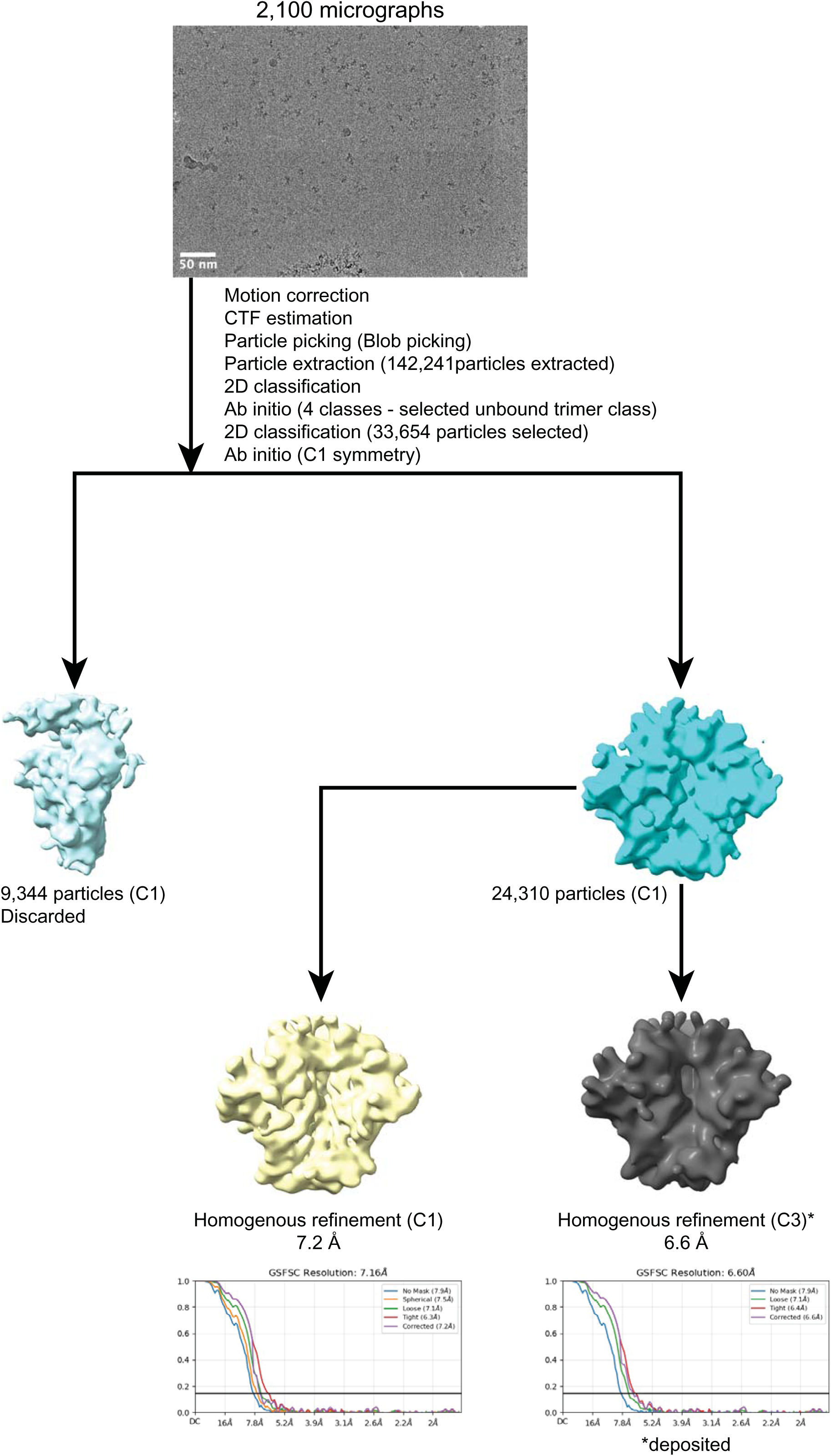
Data processing of the 3nv.2 SOSIP dataset.

**Table S1:** EM data collection and processing statistics.

| **3nv.2 SOSIP** |  |
| --- | --- |
| **Data collection conditions** |  |
| Microscope | Talos Arctica |
| Voltage (kV) | 200 |
| Camera | Gatan K3 |
| Magnification | 45,000x |
| Frames per movie | 40 |
| Recording mode | counting |
| Dose rate (e-/pixel/s) | 21.234 |
| Total electron dose (e-/Å<sup>2</sup>) | 45 |
| Defocus range (µm) | -0.8 to -1 |
| Pixel size (Å) | 0.435 (super-resolution) |
| Micrographs collected | 2,100 |
| Total extracted particles | 142,241 |
| **EMD** |  |
| Particles in class | 24,310 |
| Symmetry | C3 |
| Map resolution (Å) | 6.6 |
| FSC threshold | 0.143 |

## References

Allers, K., G. Hutter, J. Hofmann, C. Loddenkemper, K. Rieger, E. Thiel, and T. Schneider. 2011. Evidence for the cure of HIV infection by CCR5Delta32/Delta32 stem cell transplantation. Blood, 117, 2791–9.

Barnes, C. O., H. B. Gristick, N. T. Freund, A. Escolano, A. Y. Lyubimov, H. Hartweger, A. P. West, Jr., A. E. Cohen, M. C. Nussenzweig, and P. J. Bjorkman. 2018. Structural characterization of a highly-potent V3-glycan broadly neutralizing antibody bound to natively-glycosylated HIV-1 envelope. Nat Commun, 9, 1251.

Bonsignori, M., K. K. Hwang, X. Chen, C. Y. Tsao, L. Morris, E. Gray, D. J. Marshall, J. A. Crump, S. H. Kapiga, N. E. Sam, et al. 2011. Analysis of a clonal lineage of HIV-1 envelope V2/V3 conformational epitope-specific broadly neutralizing antibodies and their inferred unmutated common ancestors. J Virol, 85, 9998–10009.

Bonsignori, M., E. F. Kreider, D. Fera, R. R. Meyerhoff, T. Bradley, K. Wiehe, S. M. Alam, B. Aussedat, W. E. Walkowicz, K. K. Hwang, et al. 2017a. Staged induction of HIV-1 glycan-dependent broadly neutralizing antibodies. Sci Transl Med, 9.

Bonsignori, M., H. X. Liao, F. Gao, W. B. Williams, S. M. Alam, D. C. Montefiori, and B. F. Haynes. 2017b. Antibody-virus co-evolution in HIV infection: paths for HIV vaccine development. Immunol Rev, 275, 145–160.

Borst, A. J., C. E. Weidle, M. D. Gray, B. Frenz, J. Snijder, M. G. Joyce, I. S. Georgiev, G. B. Stewart-Jones, P. D. Kwong, A. T. Mcguire, et al. 2018. Germline VRC01 antibody recognition of a modified clade C HIV-1 envelope trimer and a glycosylated HIV-1 gp120 core. Elife, 7.

Brune, K. D., D. B. Leneghan, I. J. Brian, A. S. Ishizuka, M. F. Bachmann, S. J. Draper, S. Biswas, and M. Howarth. 2016. Plug-and-Display: decoration of Virus-Like Particles via isopeptide bonds for modular immunization. Sci Rep, 6, 19234.

Bruun, T. U. J., A. C. Andersson, S. J. Draper, and M. Howarth. 2018. Engineering a Rugged Nanoscaffold To Enhance Plug-and-Display Vaccination. ACS Nano, 12, 8855–8866.

Caniels, T. G., M. Medina-Ramirez, J. Zhang, A. Sarkar, S. Kumar, A. Labranche, R. Derking, J. D. Allen, J. L. Snitselaar, J. Capella-Pujol, et al. 2023. Germline-targeting HIV-1 Env vaccination induces VRC01-class antibodies with rare insertions. Cell Rep Med, 4, 101003.

Chen, X., T. Zhou, S. D. Schmidt, H. Duan, C. Cheng, G. Y. Chuang, Y. Gu, M. K. Louder, B. C. Lin, C. H. Shen, et al. 2021. Vaccination induces maturation in a mouse model of diverse unmutated VRC01-class precursors to HIV-neutralizing antibodies with >50% breadth. Immunity, 54, 324–339 e8.

Cohen, A. A., P. N. P. Gnanapragasam, Y. E. Lee, P. R. Hoffman, S. Ou, L. M. Kakutani, J. R. Keeffe, H. J. Wu, M. Howarth, A. P. West, et al. 2021. Mosaic nanoparticles elicit cross-reactive immune responses to zoonotic coronaviruses in mice. Science, 371, 735–741.

Cohen, A. A., J. R. Keeffe, A. Schiepers, S. E. Dross, A. J. Greaney, A. V. Rorick, H. Gao, P. N. P. Gnanapragasam, C. Fan, A. P. West, et al. 2024. Mosaic sarbecovirus nanoparticles elicit cross-reactive responses in pre-vaccinated animals. bioRxiv.

Cohen, A. A., N. Van Doremalen, A. J. Greaney, H. Andersen, A. Sharma, T. N. Starr, J. R. Keeffe, C. Fan, J. E. Schulz, P. N. P. Gnanapragasam, et al. 2022. Mosaic RBD nanoparticles protect against challenge by diverse sarbecoviruses in animal models. Science, 377, eabq0839.

Dam, K. A., C. O. Barnes, H. B. Gristick, T. Schoofs, P. N. P. Gnanapragasam, M. C. Nussenzweig, and P. J. Bjorkman. 2022a. HIV-1 CD4-binding site germline antibody-Env structures inform vaccine design. Nat Commun, 13, 6123.

Dam, K. A., P. S. Mutia, and P. J. Bjorkman. 2022b. Comparing methods for immobilizing HIV-1 SOSIPs in ELISAs that evaluate antibody binding. Sci Rep, 12, 11172.

Delaitsch, A. T., J. R. Keeffe, H. B. Gristick, J. A. Lee, W. Ding, W. Liu, A. N. Skelly, G. M. Shaw, B. H. Hahn, and P. J. Bjorkman. 2024. Neutralizing antibodies elicited in macaques recognize V3 residues on altered conformations of HIV-1 Env trimer. NPJ Vaccines, 9, 240.

Derking, R., G. Ozorowski, K. Sliepen, A. Yasmeen, A. Cupo, J. L. Torres, J. P. Julien, J. H. Lee, T. Van Montfort, S. W. De Taeye, et al. 2015. Comprehensive Antigenic Map of a Cleaved Soluble HIV-1 Envelope Trimer. PLoS Pathog, 11, e1004767.

Derking, R., and R. W. Sanders. 2021. Structure-guided envelope trimer design in HIV-1 vaccine development: a narrative review. J Int AIDS Soc, 24 Suppl 7, e25797.

Doria-Rose, N. A., C. A. Schramm, J. Gorman, P. L. Moore, J. N. Bhiman, B. J. Dekosky, M. J. Ernandes, I. S. Georgiev, H. J. Kim, M. Pancera, et al. 2014. Developmental pathway for potent V1V2-directed HIV-neutralizing antibodies. Nature, 509, 55–62.

Escolano, A., H. B. Gristick, M. E. Abernathy, J. Merkenschlager, R. Gautam, T. Y. Oliveira, J. Pai, A. P. West, Jr., C. O. Barnes, A. A. Cohen, et al. 2019. Immunization expands B cells specific to HIV-1 V3 glycan in mice and macaques. Nature, 570, 468–473.

Escolano, A., H. B. Gristick, R. Gautam, A. T. Delaitsch, M. E. Abernathy, Z. Yang, H. Wang, M. a. G. Hoffmann, Y. Nishimura, Z. Wang, et al. 2021. Sequential immunization of macaques elicits heterologous neutralizing antibodies targeting the V3-glycan patch of HIV-1 Env. Sci Transl Med, 13, eabk1533.

Escolano, A., J. M. Steichen, P. Dosenovic, D. W. Kulp, J. Golijanin, D. Sok, N. T. Freund, A. D. Gitlin, T. Oliveira, T. Araki, et al. 2016. Sequential Immunization Elicits Broadly Neutralizing Anti-HIV-1 Antibodies in Ig Knockin Mice. Cell, 166, 1445–1458 e12.

Freund, N. T., H. Wang, L. Scharf, L. Nogueira, J. A. Horwitz, Y. Bar-On, J. Golijanin, S. A. Sievers, D. Sok, H. Cai, et al. 2017. Coexistence of potent HIV-1 broadly neutralizing antibodies and antibody-sensitive viruses in a viremic controller. Sci Transl Med, 9.

Gautam, R., Y. Nishimura, A. Pegu, M. C. Nason, F. Klein, A. Gazumyan, J. Golijanin, A. Buckler-White, R. Sadjadpour, K. Wang, et al. 2016. A single injection of anti-HIV-1 antibodies protects against repeated SHIV challenges. Nature, 533, 105–9.

Gristick, H. B., H. Hartweger, M. Loewe, J. Van Schooten, V. Ramos, T. Y. Oliveira, Y. Nishimura, N. S. Koranda, A. Wall, K. H. Yao, et al. 2023. CD4 binding site immunogens elicit heterologous anti-HIV-1 neutralizing antibodies in transgenic and wild-type animals. Sci Immunol, 8, eade6364.

Gristick, H. B., L. Von Boehmer, A. P. West, Jr., M. Schamber, A. Gazumyan, J. Golijanin, M. S. Seaman, G. Fatkenheuer, F. Klein, M. C. Nussenzweig, et al. 2016. Natively glycosylated HIV-1 Env structure reveals new mode for antibody recognition of the CD4-binding site. Nat Struct Mol Biol, 23, 906–915.

Jardine, J., J. P. Julien, S. Menis, T. Ota, O. Kalyuzhniy, A. Mcguire, D. Sok, P. S. Huang, S. Macpherson, M. Jones, et al. 2013. Rational HIV immunogen design to target specific germline B cell receptors. Science, 340, 711–6.

Jardine, J. G., D. W. Kulp, C. Havenar-Daughton, A. Sarkar, B. Briney, D. Sok, F. Sesterhenn, J. Ereno-Orbea, O. Kalyuzhniy, I. Deresa, et al. 2016. HIV-1 broadly neutralizing antibody precursor B cells revealed by germline-targeting immunogen. Science, 351, 1458–63.

Jardine, J. G., T. Ota, D. Sok, M. Pauthner, D. W. Kulp, O. Kalyuzhniy, P. D. Skog, T. C. Thinnes, D. Bhullar, B. Briney, et al. 2015. HIV-1 VACCINES. Priming a broadly neutralizing antibody response to HIV-1 using a germline-targeting immunogen. Science, 349, 156–61.

Joyce, M. G., I. S. Georgiev, Y. Yang, A. Druz, H. Geng, G. Y. Chuang, Y. D. Kwon, M. Pancera, R. Rawi, M. Sastry, et al. 2017. Soluble Prefusion Closed DS-SOSIP.664-Env Trimers of Diverse HIV-1 Strains. Cell Rep, 21, 2992–3002.

Keeble, A. H., P. Turkki, S. Stokes, I. N. A. Khairil Anuar, R. Rahikainen, V. P. Hytönen, and M. Howarth. 2019. Approaching infinite affinity through engineering of peptide–protein interaction. Proceedings of the National Academy of Sciences, 116, 26523–26533.

Kong, R., H. Duan, Z. Sheng, K. Xu, P. Acharya, X. Chen, C. Cheng, A. S. Dingens, J. Gorman, M. Sastry, et al. 2019. Antibody Lineages with Vaccine-Induced Antigen-Binding Hotspots Develop Broad HIV Neutralization. Cell, 178, 567–584.e19.

Kong, R., K. Xu, T. Zhou, P. Acharya, T. Lemmin, K. Liu, G. Ozorowski, C. Soto, J. D. Taft, R. T. Bailer, et al. 2016. Fusion peptide of HIV-1 as a site of vulnerability to neutralizing antibody. Science, 352, 828–33.

Lee, J. H., G. Ozorowski, and A. B. Ward. 2016. Cryo-EM structure of a native, fully glycosylated, cleaved HIV-1 envelope trimer. Science, 351, 1043–8.

Li, H., S. Wang, R. Kong, W. Ding, F. H. Lee, Z. Parker, E. Kim, G. H. Learn, P. Hahn, B. Policicchio, et al. 2016. Envelope residue 375 substitutions in simian-human immunodeficiency viruses enhance CD4 binding and replication in rhesus macaques. Proc Natl Acad Sci U S A, 113, E3413–22.

Li, W., Z. Qin, E. Nand, M. W. Grunst, J. R. Grover, J. W. Bess, Jr., J. D. Lifson, M. B. Zwick, H. D. Tagare, P. D. Uchil, et al. 2023. HIV-1 Env trimers asymmetrically engage CD4 receptors in membranes. Nature, 623, 1026–1033.

Li, Z., W. Li, M. Lu, J. Bess, C. W. Chao, J. Gorman, D. S. Terry, B. Zhang, T. Zhou, S. C. Blanchard, et al. 2020. Subnanometer structures of HIV-1 envelope trimers on aldrithiol-2-inactivated virus particles. Nature Structural & Molecular Biology, 27, 726–734.

López-Sagaseta, J., E. Malito, R. Rappuoli, and M. J. Bottomley. 2016. Self-assembling protein nanoparticles in the design of vaccines. Computational and Structural Biotechnology Journal, 14, 58–68.

Mastronarde, D. N. 2005. Automated electron microscope tomography using robust prediction of specimen movements. J Struct Biol, 152, 36–51.

Mccoy, L. E., and D. R. Burton. 2017. Identification and specificity of broadly neutralizing antibodies against HIV. Immunol Rev, 275, 11–20.

Mcguire, A. T., M. D. Gray, P. Dosenovic, A. D. Gitlin, N. T. Freund, J. Petersen, C. Correnti, W. Johnsen, R. Kegel, A. B. Stuart, et al. 2016. Specifically designed immunogens select and activate B cells expressing precursors of broadly neutralizing human antibodies to HIV-1 in knock-in mice. Nature Comm, 7, 10618.

Mcguire, A. T., S. Hoot, A. M. Dreyer, A. Lippy, A. Stuart, K. W. Cohen, J. Jardine, S. Menis, J. F. Scheid, A. P. West, et al. 2013. Engineering HIV envelope protein to activate germline B cell receptors of broadly neutralizing anti-CD4 binding site antibodies. J Exp Med, 210, 655–63.

Medina-Ramirez, M., F. Garces, A. Escolano, P. Skog, S. W. De Taeye, I. Del Moral-Sanchez, A. T. Mcguire, A. Yasmeen, A. J. Behrens, G. Ozorowski, et al. 2017. Design and crystal structure of a native-like HIV-1 envelope trimer that engages multiple broadly neutralizing antibody precursors in vivo. J Exp Med, 214, 2573–2590.

Mkhize, N. N., A. E. J. Yssel, H. Kaldine, R. T. Van Dorsten, A. S. Woodward Davis, N. Beaume, D. Matten, B. Lambson, T. Modise, P. Kgagudi, et al. 2023. Neutralization profiles of HIV-1 viruses from the VRC01 Antibody Mediated Prevention (AMP) trials. PLoS Pathog, 19, e1011469.

Molinos-Albert, L. M., E. Baquero, M. Bouvin-Pley, V. Lorin, C. Charre, C. Planchais, J. D. Dimitrov, V. Monceaux, M. Vos, A. V. S. Group, et al. 2023. Anti-V1/V3-glycan broadly HIV-1 neutralizing antibodies in a post-treatment controller. Cell Host Microbe, 31, 1275–1287 e8.

Montefiori, D. C. 2005. Evaluating neutralizing antibodies against HIV, SIV, and SHIV in luciferase reporter gene assays. *Curr Protoc Immunol*, Chapter 12, Unit 12 11.

Pettersen, E. F., T. D. Goddard, C. C. Huang, E. C. Meng, G. S. Couch, T. I. Croll, J. H. Morris, and T. E. Ferrin. 2021. UCSF ChimeraX: Structure visualization for researchers, educators, and developers. Protein Sci, 30, 70–82.

Punjani, A., J. L. Rubinstein, D. J. Fleet, and M. A. Brubaker. 2017. cryoSPARC: algorithms for rapid unsupervised cryo-EM structure determination. Nat Methods, 14, 290–296.

Rahikainen, R., P. Rijal, T. K. Tan, H. J. Wu, A. C. Andersson, J. R. Barrett, T. A. Bowden, S. J. Draper, A. R. Townsend, and M. Howarth. 2021. Overcoming Symmetry Mismatch in Vaccine Nanoassembly through Spontaneous Amidation. Angew Chem Int Ed Engl, 60, 321–330.

Rich, R. L., and D. G. Myszka. 2010. Grading the commercial optical biosensor literature-Class of 2008: ‘The Mighty Binders’. J Mol Recognit, 23, 1–64.

Rich, R. L., and D. G. Myszka. 2011. Survey of the 2009 commercial optical biosensor literature. J Mol Recognit, 24, 892–914.

Roark, R. S., H. Li, W. B. Williams, H. Chug, R. D. Mason, J. Gorman, S. Wang, F.-H. Lee, J. Rando, M. Bonsignori, et al. 2021. Recapitulation of HIV-1 Env-antibody coevolution in macaques leading to neutralization breadth. Science, 371, eabd2638.

Sanders, R. W., R. Derking, A. Cupo, J.-P. Julien, A. Yasmeen, N. De Val, H. J. Kim, C. Blattner, A. T. De La Peña, J. Korzun, et al. 2013. A Next-Generation Cleaved, Soluble HIV-1 Env Trimer, BG505 SOSIP.664 gp140, Expresses Multiple Epitopes for Broadly Neutralizing but Not Non-Neutralizing Antibodies. PLoS Pathogens, 9, e1003618–20.

Saunders, K. O., K. Wiehe, M. Tian, P. Acharya, T. Bradley, S. M. Alam, E. P. Go, R. Scearce, L. Sutherland, R. Henderson, et al. 2019. Targeted selection of HIV-specific antibody mutations by engineering B cell maturation. Science, 366.

Shingai, M., O. K. Donau, R. J. Plishka, A. Buckler-White, J. R. Mascola, G. J. Nabel, M. C. Nason, D. Montefiori, B. Moldt, P. Poignard, et al. 2014. Passive transfer of modest titers of potent and broadly neutralizing anti-HIV monoclonal antibodies block SHIV infection in macaques. J Exp Med, 211, 2061–74.

Silva, M., Y. Kato, M. B. Melo, I. Phung, B. L. Freeman, Z. Li, K. Roh, J. W. Van Wijnbergen, H. Watkins, C. A. Enemuo, et al. 2021. A particulate saponin/TLR agonist vaccine adjuvant alters lymph flow and modulates adaptive immunity. Sci Immunol, 6, eabf1152.

Simonich, C. A., K. L. Williams, H. P. Verkerke, J. A. Williams, R. Nduati, K. K. Lee, and J. Overbaugh. 2016. HIV-1 Neutralizing Antibodies with Limited Hypermutation from an Infant. Cell, 166, 77–87.

Sliepen, K., M. Medina-Ramirez, A. Yasmeen, J. P. Moore, P. J. Klasse, and R. W. Sanders. 2015. Binding of inferred germline precursors of broadly neutralizing HIV-1 antibodies to native-like envelope trimers. Virology, 486, 116–20.

Slifka, M. K., and I. J. Amanna. 2019. Role of Multivalency and Antigenic Threshold in Generating Protective Antibody Responses. Frontiers in Immunology, 10.

Sok, D., M. J. Van Gils, M. Pauthner, J. P. Julien, K. L. Saye-Francisco, J. Hsueh, B. Briney, J. H. Lee, K. M. Le, P. S. Lee, et al. 2014. Recombinant HIV envelope trimer selects for quaternary-dependent antibodies targeting the trimer apex. Proc Natl Acad Sci U S A, 111, 17624–9.

Stamatatos, L., M. Pancera, and A. T. Mcguire. 2017. Germline-targeting immunogens. Immunol Rev, 275, 203–216.

Steichen, J. M., D. W. Kulp, T. Tokatlian, A. Escolano, P. Dosenovic, R. L. Stanfield, L. E. Mccoy, G. Ozorowski, X. Hu, O. Kalyuzhniy, et al. 2016. HIV Vaccine Design to Target Germline Precursors of Glycan-Dependent Broadly Neutralizing Antibodies. Immunity, 45, 483–96.

Steichen, J. M., Y. C. Lin, C. Havenar-Daughton, S. Pecetta, G. Ozorowski, J. R. Willis, L. Toy, D. Sok, A. Liguori, S. Kratochvil, et al. 2019. A generalized HIV vaccine design strategy for priming of broadly neutralizing antibody responses. Science, 366.

Turner, H. L., A. M. Jackson, S. T. Richey, L. M. Sewall, A. Antanasijevic, L. Hangartner, and A. B. Ward. 2023. Protocol for analyzing antibody responses to glycoprotein antigens using electron-microscopy-based polyclonal epitope mapping. STAR Protoc, 4, 102476.

Walker, L. M., S. K. Phogat, P. Y. Chan-Hui, D. Wagner, P. Phung, J. L. Goss, T. Wrin, M. D. Simek, S. Fling, J. L. Mitcham, et al. 2009. Broad and Potent Neutralizing Antibodies from an African Donor Reveal a New HIV-1 Vaccine Target. Science, 326, 285–289.

Walsh, S. R., and M. S. Seaman. 2021. Broadly Neutralizing Antibodies for HIV-1 Prevention. Front Immunol, 12, 712122.

Wang, H., C. Cheng, J. L. Dal Santo, C. Shen, T. Bylund, A. R. Henry, C. A. Howe, J. Hwang, N. C. Morano, D. J. Morris, et al. 2024. Potent cross-reactive HIV-1 neutralization in fusion peptide-primed SHIV-infected macaques. Cell, in press.

Ward, A. B., and I. A. Wilson. 2017. The HIV-1 envelope glycoprotein structure: nailing down a moving target. Immunol Rev, 275, 21–32.

Willis, J. R., Z. T. Berndsen, K. M. Ma, J. M. Steichen, T. Schiffner, E. Landais, A. Liguori, O. Kalyuzhniy, J. D. Allen, S. Baboo, et al. 2022. Human immunoglobulin repertoire analysis guides design of vaccine priming immunogens targeting HIV V2-apex broadly neutralizing antibody precursors. Immunity, 55, 2149–2167 e9.

Wyatt, R., and J. Sodroski. 1998. The HIV-1 envelope glycoproteins: fusogens, antigens, and immunogens. Science, 280, 1884–8.

Xiao, X., W. Chen, Y. Feng, Z. Zhu, P. Prabakaran, Y. Wang, M. Y. Zhang, N. S. Longo, and D. S. Dimitrov. 2009. Germline-like predecessors of broadly neutralizing antibodies lack measurable binding to HIV-1 envelope glycoproteins: implications for evasion of immune responses and design of vaccine immunogens. Biochem Biophys Res Commun, 390, 404–9.

Zhang, P., E. Narayanan, Q. Liu, Y. Tsybovsky, K. Boswell, S. Ding, Z. Hu, D. Follmann, Y. Lin, H. Miao, et al. 2021. A multiclade env-gag VLP mRNA vaccine elicits tier-2 HIV-1-neutralizing antibodies and reduces the risk of heterologous SHIV infection in macaques. Nat Med, 27, 2234–2245.

Zhou, T., R. M. Lynch, L. Chen, P. Acharya, X. Wu, N. A. Doria-Rose, M. G. Joyce, D. Lingwood, C. Soto, R. T. Bailer, et al. 2015. Structural Repertoire of HIV-1-Neutralizing Antibodies Targeting the CD4 Supersite in 14 Donors. Cell, 161, 1280–92.

